# Human RNA ligase 1 as a novel regulator of ribosome function and translation under oxidative stress

**DOI:** 10.1101/2025.10.23.684095

**Authors:** Florian Michael Stumpf, Marissa Glauner, Jasmin Jansen, Florian Stengel, Andreas Marx

## Abstract

Human RNA ligase 1 (Rlig1) is a recently identified human 5’-3’ RNA ligase essential for maintaining 28S rRNA integrity and promoting cell survival under oxidative stress. Although the enzymatic activity of Rlig1 implies a role in RNA maintenance or repair, its broader molecular context remains insufficiently characterised. Here we identify cellular interactors of Rlig1 using affinity enrichment coupled to mass spectrometry in a HEK293 Rlig1-KO model. Our approach revealed several RNA and DNA surveillance, degradation and repair proteins as well as a diverse set of RNA binding and RNA processing enzymes as potential interactors. Notably, ribosomal proteins were highly enriched, suggesting a link between Rlig1 and the translational machinery. Cross-linking coupled to mass spectrometry confirmed that Rlig1 interacts with 80S ribosomes in vitro and preferentially binds the large ribosomal subunit near the 28S rRNA. Polysome profiling corroborated this interaction, as Rlig1 was recruited to ribosomal fractions containing the large ribosomal subunit in response to oxidative stress. Moreover, Rlig1-deficient cells exhibited earlier polysome loss and significantly impaired translational activity compared to their WT counterparts under these conditions. These results highlight a potential novel role for Rlig1 in safeguarding translation during oxidative stress.

**GRAPHICAL ABSTRACT:** 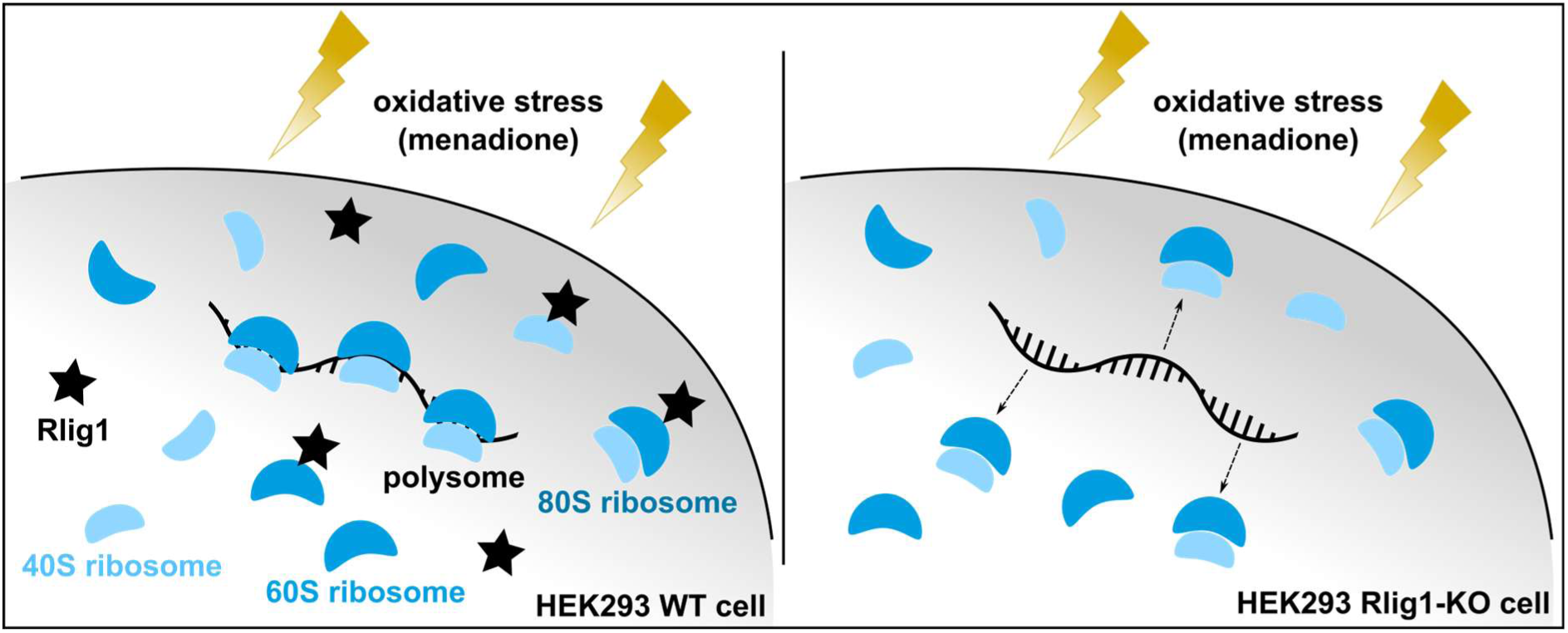

## INTRODUCTION

DNA repair has been extensively studied over the past decades, significantly advancing our understanding of genome maintenance. By now, a large number of DNA repair pathways have been described that guarantee the integrity of the genome.(1–3) In contrast, RNA repair remains poorly characterized. The prevailing view holds that damaged RNA is primarily eliminated through degradation, followed by resynthesis from an intact genomic template.(4, 5) The only two mechanisms of RNA repair known to date are the reversal of methylations, such as 1-methyl adenosine (1meA) and 3-methyl cytosine (3meC) by the AlkB family(6) and the removal of RNA-protein crosslinks.(7–9) However, not only DNA but also specific RNA types have been shown to exist and persist in cells for longer periods of time.(10–12) It was also shown that RNA is be subject to certain types of cellular damages at an even greater rate than DNA.(13–15) Moreover, RNA is estimated to be ∼4-fold more abundant than DNA in mammalian cells.(16) Thus, repairing damaged RNA, rather than degrading and resynthesizing it, could be energetically favourable for the cell, particularly in the case of long-lived RNAs.(9)

The recent discovery of the 5’-3’ human RNA ligase Rlig1 supports the assumption that additional, yet unidentified RNA repair mechanisms could exist in human cells. (17)

Rlig1 is the first described human 5’-3’ RNA ligase, able to fuse RNA strands with 5’-PO_4_ ends with 3’-OH ends.(17) As a member of the nucleotidyltransferase family, characterized by its conserved sequence motif KxxG in the active centre, the ligation mechanism of Rlig1 operates via a three-step mechanism.(17–19) Rlig1 is able to autoAMPylate itself at the position of lysine K57 located within its active site. Subsequently, Rlig1 transfers the AMP-moiety onto the 5’-PO_4_ of the first RNA strand that is to be ligated. This is followed by a subsequent nucleophilic attack by the 3’-OH of the second RNA strand on the formed RNA-adenylate resulting in the ligated RNA product.

We previously reported that Rlig1-deficient HEK293 cells exhibited increased degradation of 28S rRNA and reduced viability following menadione-induced oxidative stress, compared to wild-type (WT) cells.(17) Similar effects were observed upon chemical inhibition of Rlig1.(20) Further research demonstrated that, in addition to menadione, a small number of other specific compounds were also capable of reducing the viability of HEK293 Rlig1-knockout (KO) cells compared to WT cells.(21) Consequently, Rlig1-deficient HEK293 cells proved to be more susceptible to certain cellular stressors than HEK293 WT cells, supporting the hypothesis of a role for Rlig1 in RNA maintenance or repair. Moreover, recent studies reported altered tRNA, snoRNA, and snRNA levels in the brains of Rlig1-KO mice, (22) showed that Rlig1 is integral for circRNA biogenesis during viral infections(23) and that it promotes proliferation of triple-negative breast cancer cells by interacting with ERK, highlighting its potential oncogenic role.(24)

To gain insight into the cellular roles of Rlig1, we sought to elucidate its interactome. To identify potential cellular interactors of Rlig1 in human cells, we employed affinity enrichment-mass spectrometry using recombinantly expressed active (AMP-Rlig1) and inactive (K57A-Rlig1) variants immobilised on Strep-Tactin beads, which were then incubated with lysates from an HEK293 Rlig1-KO cell line. This approach identified RNA-binding and processing enzymes as the most prominent candidates for Rlig1 interaction, notably some of which are involved in RNA surveillance and degradation. For instance, ANGEL2 and CLP1, along with members of the short-patch base excision repair pathway of DNA were enriched in our experiments. This is of particular interest as these proteins had previously been hypothesised to be involved in RNA maintenance or repair.(17, 25–27)

Interestingly, ribosomal proteins were also significantly enriched. Subsequent experiments revealed that Rlig1 interacts with the ribosome and is recruited to it under oxidative stress induced by menadione. Menadione treatment also reduced translational activity in Rlig1-KO cells relative to WT cells, evidenced by fewer polysomes. This suggests an important role of Rlig1 in the maintenance of translational processes under oxidative stress and supports the hypothesis that Rlig1 is involved in the quality control or repair of RNA under these conditions.

## MATERIAL AND METHODS

### General procedures

#### Cell culture

HEK293 WT and Rlig1-KO cells were cultured in DMEM Glutamax + 10% FCS in 15 cm standard cell culture plates (Sarstedt). The cells were passaged every other day, and the cells were incubated at 37°C, 7.5% CO_2_ and 95% humidity.

## SDS-PAGE

For the analysis of protein samples via SDS-PAGE, 6x SDS loading dye was added to the samples to reach a final concentration of 1x. Subsequently, the samples were heated and denatured (95°C, 5 min). The proteins were separated utilising a PAGE gel consisting of a stacking gel (5 % acrylamide) and a resolving gel (12.5 % acrylamide). Gel electrophoresis was performed with a BioRad system in 1x SDS running buffer (25 mM Tris-HCl, 200 mM glycine, 0.1 % (w/v) SDS) at 35 mA for one gel or 65 mA for two gels until the desired separation was achieved. As marker, a prestained protein ladder (Thermo Scientific) was used. The resulting gel with the separated proteins was subsequently stained with Krypton™ Fluorescence Protein Stain (Thermo Scientific) according to the manufacturer’s instructions or transferred on a PVDF membrane (0.2 µm, GE Healthcare) via western blot transfer.

### Western blot

The proteins were separated via SDS-PAGE as described above. The western blot transfer of proteins from the gel onto a PVDF membrane (0.2 µm, GE Healthcare) was performed in 1x Western blot transfer buffer (25 mM Tris-HCl pH 8.3, 100 mM glycine) for 90 min at 60 V. After the transfer, the membrane was blocked with 5 % milk in PBS-T for 1 h. Then the membrane was incubated with the respective primary antibody (Rlig1 (sc-390730, Santa Cruz Biotechnology), RPL10L (ABC-AP17603a, Abcepta), RPS20 (ab133776, Abcam) or puromycin (MABE343, Merck)) according to the manufacturer’s instructions. After washing the membrane (3 x 10 min, PBS-T), the suiting secondary antibody (1:20’000 in PBS-T, α-mouse or α-rabbit) was added to the membrane and incubated for 1 h. Afterwards, the membrane was washed again (3 x 10 min, PBS-T) and incubated with Pierce™ ECL Western Blotting Substrate (Thermo Scientific) or SuperSignal™ West Femto Maximum Sensitivity Substrate (Thermo Scientific). Then, the membrane was subjected to chemiluminescence-imaging with an Amersham imager 600 (GE Healthcare).

### BCA determination of protein concentration

For the determination of protein concentration in a sample, a Bicinchoninic acid assay (BCA), the Pierce™ BCA Protein Assay Kit (Thermo Scientific) was utilised according to the manufacturers protocol. The calibration was achieved by diluting BSA in the same buffer as the samples in a range of 0.125 mg/mL to 2.000 mg/mL. For the assay, 10 µL of each sample or standard was mixed with 200 µL of the provided BCA solution in a 96-well plate. The mixture was incubated at 37°C for 30 min in the dark and subsequently read out with a Victor^3^ 1420 Multilabel Counter (PerkinElmer).

### Elucidation of protein-protein interaction partners of Rlig1

#### Expression and purification of recombinant Rlig1 variants

The Rlig1 variants were expressed as previously reported.(17)

### Cell pellet production

For the cell pellets, 8 x 10^6^ cells were seeded in 20 mL medium in 15 cm cell culture plates. After 48 h at approximately 90 % confluency, the cells were harvested. For stressed cells, the medium was removed and 10 mL new medium containing 40 µM menadione in EtOH (0.1 %) was added and incubated (1 h, 37°C). Afterwards the medium was removed and 5 mL ice-cold DPBS was added to the cells. For unstressed cells the incubation with menadione was omitted and 5 mL ice-cold DPBS was directly added after removing the medium. The cells were harvested with a cell scraper and centrifuged (10 min, 500 g, 4°C). The resulting pellets were resuspended in 400 µL ice-cold DPBS and the cells of four plates were combined and centrifuged (10 min, 500 g, 4°C). After removing the DPBS the cell pellet was stored frozen (-80°C).

### Lysate preparation for affinity enrichment

Directly before the experiment the frozen cell pellet was lysed by resuspending it in 750 µL ice-cold lysis buffer (25 mM Tris pH 7.5, 150 mM NaCl, 1 mM EDTA, 1 % (v/v) NP40, 1 mM DTT, 0.1 M Pefabloc® SC, 1 µg/mL aproptinin/leupeptin) and incubation on ice for 20 min. After the incubation, the lysate was cleared by centrifugation in an Eppendorf 5430R centrifuge (22’000 g, 30 min, 4°C). The protein concentration of the supernatant was determined by BCA as described above and adjusted to 6.67 mg/mL with lysis buffer. Subsequently, RNase If (6000 U/mL in lysis buffer) was added to the cell lysate (end concentration 600 U/mL with 6 mg/mL protein concentration) and incubated in a thermomixer (15 min, 37°C, 850 rpm) to result in the RNA digested cell lysate. The RNA digestion was controlled via a 4150 TapeStation system (Agilent).

### Tapestation control of the RNA digested cell lysate

250 µL TRIzol™ reagent was given to 25 µL cell lysate in a 500 µL low-binding reaction tube. The mixture was vortexed (1 min) and incubated at RT (5 min). Afterwards, chloroform (250 µL) was added and the mixture was again vortexed (1 min) and incubated at RT (5 min). The sample was centrifuged (12’000 g, 15 min, 4°C) and the upper aqueous phase was transferred into a new 500 µL reaction tube. To this aqueous phase chloroform (62.5 µL) was added and the mixture was vortexed (1 min) and incubated at RT (5 min) before being centrifuged again (12’000 g, 15 min, 4°C). The upper aqueous phase was transferred into a new 500 µL reaction tube and ice-cold isopropanol (170 µL)was added. The mixture was vortexed (1 min), flash frozen in liquid nitrogen and incubated (-80°C, 18 h). The sample was carefully thawed and centrifuged (20’700 g, 30 min, 4°C) and the supernatant was discarded. To the transparent pellet ice-cold 80 % EtOH (250 µL) was added and the sample was vortexed (1 min) and centrifuged again (20’700 g, 30 min, 4°C). After discarding the supernatant, the white RNA pellet was air dried (10 min) and redissolved in MQ water (40 µL). The resulting sample was analysed via an HSRNA ScreenTape following the manufacturer’s protocol on a 4150 TapeStation system (Agilent).

### Affinity enrichment using Rlig1 as bait

For the affinity enrichment of Rlig1, Strep-Tactin® Superflow® bead slurry (50 µL, IBA Lifesciences) was prepared in 500 µL low-binding reaction tubes. The beads were equilibrated (3 x 400 µL) with washing buffer (25 mM Tris pH 7.5, 50 mM NaCl, 1 mM DTT) and kept on ice. His_6_-Strep-AMP-Rlig1 or His_6_-Strep-K57A-Rlig1 (100 µL, 0.5 mg/mL in washing buffer) was added to the equilibrated beads (50 µL). Washing buffer (100 µL) was used as negative control.

The bead suspension was incubated in an overhead shaker (60 min, 4°C). Washing buffer (400 µL) was added to the beads and the sample was transferred to Mobicol spin columns (MoBiTec). The reaction tube that contained the beads was rinsed one time with washing buffer (100 µL) and the bead containing solutions were combined. The beads were washed (4 x 500 µL, washing buffer) and the RNA digested cell lysate (250 µL, 6 mg/mL) was added to the prepared beads and incubated on an overhead shaker (20 h, 4°C). The beads were washed (5 x 500 µL, washing buffer) and the protein was eluted with elution buffer (50 mM NH_4_HCO_3_, 0.8 mM biotin) (4 x 100 µL, 10 min, 37°C, 800 rpm). The eluate was dried in a SpeedVac (2 h, vacuum, no heating) and stored at -20°C.

### Tryptic In-solution digest

The dried samples were dissolved in urea (100 µL, 8 M). Tris(2-carboxyethyl)phosphin (5 mM) was added and the samples were incubated (30 min, 37°C, 600 rpm). Afterwards, chloroacetamide (10 mM) was added to the solution and incubated (30 min, 23°C, 600 rpm, in the dark). The solution was diluted with NH_4_HCO_3_ (700 µL, 50 mM) and the pH was controlled to be between pH 7-8. Sequencing grade modified trypsin (2.5 µg, Promega (cat. nr. V5111)) was added, and the sample was incubated (20 h, 37°C, 600 rpm). TFA in H_2_O (10% (v/v), 5 µL, LC-MS grade) was added and the sample was freeze-dried.

### Desalting of the peptides

The freeze-dried peptides were redissolved (0.1 % TFA in 5 % ACN, 40 µL) and 10 % TFA (10 µL) was added. The pH was checked to be pH < 4. The spin tips were wetted (0.1 % TFA in 80 % ACN 20 µL) and equilibrated (0.1 % TFA, 3 x 20 µL). The sample was loaded and the flow-through was loaded a second time for better retention. The tip was washed two times (0.1 % TFA, 20 µL) and the bound peptides were eluted (0.1 % FA in 80 % ACN, 3 x 20 µL). The combined elution was dried in a SpeedVac (45 min, vacuum, no heat).

### Affinity enrichment mass spectrometry

For MS measurement the samples were redissolved (0.1 % FA, 15 µL). The desalted peptide samples were analyzed on an EASY-nLC 1200 UHPLC system (Thermo Scientific) coupled to a Q-Exactive HF mass spectrometer (Thermo Scientific) and separated on a C18 Acclaim PepMap RSLC column (50 µm x 15 cm, 2 µm, 100 Å, Thermo Scientific) with a flow rate of 300 nL/min over a 120 min gradient. After starting for 4 min with 94 % buffer A (0.1 % FA) and 6 % buffer B (80 % ACN + 0.1 % FA) the concentration of buffer B was increased to 44 % over 105 min, then set to 56 % over 5 min and finally increased to 100 % buffer B over 1 min, followed by washing and re-equilibration of the column for 5 min. The samples were measured in data independent acquisition (DIA) mode using a total of 22 isolation windows of variable window size. The isolation width and window positions were adapted from Bruderer et al.(28) MS1 scans were acquired between 300 and 1650 m/z with a resolution of 120’000 at 200 m/z. A maximum injection time of 60 ms and an AGC target of 3e6 was used. After one MS1 scan the mass spectrometer cycled once through all DIA windows. Isolated ions were fragmented via higher-energy collisional dissociation (HCD) with stepped normalized collision energies of 25.5, 27 and 30 eV and fragment ion spectra were acquired with automatic injection time at a resolution of 30’000 at 200 m/z and an AGC target of 3e6. Spectra were recorded with a fixed first mass of 200 m/z.

### Data analysis of affinity enrichment

The raw data from the MS measurements were first analysed with Spectronaut V16. Here, identification and label-free quantification of proteins was performed in library-free directDIA mode. The default settings were used with a minimum peptide length of 5 AAs. “Uniprot_sprot_2022_01_07_HUMAN.fasta” (downloaded 07/01/2022) and “Frankenfield2022_Universal_Contaminants.fasta” were used as databases. The results were exported from Spectronaut and all further analyses of the raw LFQ data was performed with Perseus V1.6.15.0. Contaminants were filtered out using the contamination database and the LFQ intensities were log_2_(x) transformed. Only proteins that were consistently identified with a minimum of three precursor peptides in all three replicates of at least one condition were subjected to further evaluation. Missing values were imputed from a normal distribution (width = 0.3, shift = 1.8) and an ANOVA multiple sample test was conducted (S_0_ = 0.1, FDR = 0.01), followed by a Post hoc Tukey’s test (FDR = 0.01). Proteins enriched solely for the bead control according to the Post hoc test were filtered out before further processing. Intenstities were normalised by z-scoring and the mean average of the triplicate measurements for each bait/condition was calculated. Only proteins with a minimum Z-score of >0.5 for the AMP-Rlig1 bait and a Z-score <0 for the bead control after the ANOVA analysis were considered as enriched for AMP-Rlig1 and considered for further analysis. GO-term analysis was performed with the DAVID tool.(29) The mass spectrometry proteomics data have been deposited to the ProteomeXchange Consortium via the PRIDE partner repository(30) with the dataset identifier PXD067856 (Username; Password: 5cD8925AZ6ug).

### Ribosomal experiments

#### Ribosome purification

For the purification of ribosomes, 5 ml HEK293 cell pellet was resuspended in 25 ml lysis buffer (50 mM HEPES pH 7.4, 300 mM NaCl, 6 mM MgCl_2_, 0.5% NP40, 20 µM phenanthrolin, 1 mM PMSF, 2 mM DTT, 1x cOmplete™ EDTA-free Protease inhibitor (Roche)) and incubated (30 min, 4°C, 400 rpm). After incubation with puromycin (1 mM, 30 min, 4°C) the lysate was cleared by two centrifugation steps (12’000 g, 10 min, 4°C). The cleared lysate was loaded on a 60% sucrose cushion (60% sucrose in 50 mM HEPES pH 7.4, 50 mM KCl, 10 mM MgCl_2_, 5 mM EDTA, 1x cOmplete™ EDTA-free Protease inhibitor (Roche)). For 8 ml of the cleared lysate 12 ml sucrose cushion was used. Afterwards, the ribosomes were pelleted by centrifugation (184’000 g, 20 h, 4°C). The 80S ribosome pellet was resuspended (50 mM HEPES pH 7.4, 300 mM KCl, 1 mM DTT) and the concentration for the 80S ribosomal fraction in µM was determined by multiplying the A260 absorbance measured via Nanodrop ND-1000 (Thermo Fisher) with the factor 0.02.

#### Ribosome sedimentation

80S ribosomes and Rlig1 were mixed in different concentrations in the reaction buffer (30 mM HEPES pH 7.4, 100 mM KAc, 5 mM MgCl_2_) in a total volume of 35 µL and incubated (15 min, 400 rpm, 12°C). Afterwards, 30 µL of the sample was loaded onto a sucrose cushion (150 µL, 25 % sucrose) in polycarbonate centrifuge tubes (Beckman Coulter, 7 x 20 mm) and centrifuged (90 min, 220’000 g, 4°C). After removing the sucrose, the resulting ribosomal pellet was carefully resuspended in 1x Laemmli sample buffer (30 µL, 30 min, 23°C, 600 rpm). Afterwards more 1x Laemmli sample buffer was added (30 µL) and the sample was denatured (5 min, 95°C). For the samples before the sedimentation, the remaining 5 µL reaction solution were diluted with 2x Laemmli sample buffer (5 µL) and the sample was denatured (5 min, 95°C). The samples were analysed by PAGE and western blot.

#### Crosslinking experiment

17, 5 µM Rlig1-AMP or apo-Rlig1 K57A was gently mixed with 67 nM 80S ribosomes in cross-linking buffer (50 mM HEPES pH 7.50, 5 mM MgCl_2_, 100 mM KCl). The mixture was incubated without agitation to allow complex formation (30 min, 37°C). Afterwards, 1.5 mM BS3-H12/D12 crosslinker (Creative Molecules Inc.) was added and the reaction mixture was incubated with gentle agitation (30 min, 37°C, 350 rpm). After the crosslinking, unreacted linker was quenched by the addition of NH_4_HCO_3_ (50 mM, 15 min). The samples were dried and subsequently dissolved in 8 M urea. The sample was reduced with Tris(2-carboxyethyl)phosphine (2.5 mM, 30 min, 37°C), alkylated with iodoacetamide (5 mM, 30 min, RT, in the dark) and digested with trypsin (Promega, 1:50 enzyme-to-substrate, 16 h, 37°C). The peptides were desalted (C18 Sep-Pak cartridges (Waters)) and crosslinked peptides were enriched by size exclusion chromatography using an ÄKTAmicro chromatography system (GE Healthcare) equipped with a SuperdexTM Peptide 3.2/30 column. Fractions containing peptides were collected and combined to four fractions. LC-MS^2^ analysis was performed on an Orbitrap Tribrid Fusion mass spectrometer coupled to an EASY-nLC 1200 system (Thermo Scientific). Peptides were separated across a 75 min gradient on an Acclaim PepMap column (150x0.05 mm, 2 µm, Thermo Scientific, P/N 164943). MS measurement was performed in data-dependent mode with a maximum cycle time of 3 sec. Full scans were acquired in the Orbitrap at a resolution of 120’000, in the scan range of 400-1500 m/z, AGC target 2e5 and a maximum injection time of 50 ms. For precursor selection, monoisotopic precursor selection was set to peptides, dynamic exclusion to 60 sec and only precursor ions with charge states 3-8 were isolated for MS/MS acquisition. Precursor ions were fragmented with 35 % CID activation and MS/MS scan was acquired in the iontrap in rapid scan mode. Crosslink analysis was done with xQuest/xProphet(31) in ion-tag mode with a precursor mass tolerance of 10 ppm. For matching of fragment ions, tolerances of 0.2 Da for common ions and 0.3 Da for crosslinked ions were applied. The database was compiled of the Rlig1 protein sequence as well as proteins that are part of the mature cytosolic ribosome. An FDR was estimated by xProphet and only hits with an FDR below 0.01 were used. The experiment was performed as biological triplicate with two technical replicates each. Crosslinks identified with ld-score ≥ 25 and deltaS < 0.95 were retained for further analysis and interlinks were filtered with the mi-filter.(32) The crosslinking data was visualized by xiNET.(33) The XL-MS data have been deposited to the ProteomeXchange Consortium via the PRIDE partner repository(30) with the dataset identifier PXD067615 (Username; Password: 7nHY32A9Vzj6).

### Polysome profiling

#### Preparation of the sucrose gradient

Sucrose solution (5% and 50% (w/v)) in polysome profiling buffer (20 mM Tris pH 7.4, 100 mM KCl, 10 mM MgCl_2_; complete EDTA-free protease inhibitor cocktail (Roche), 100 µg/mL cycloheximide, 2 mM DTT, 50 U/mL RNasin (Promega)) was prepared. A linear gradient was generated in ultracentrifuge tubes (14 x 89 mm, PC, Seton Scientific) using a Gradient Master device (BioComp) running the ’SW41 Short Gradient’ program.

#### Polysome isolation and fractionation

Two days prior the experiment, 6 x 10^6^ HEK293 WT or Rlig1-KO cells were seeded in 15 cm cell culture dishes as described in the cell culture section above. At 80% confluency, the cells were treated with menadione (40 µM or 60 µM) for oxidative stress induction for 1 h. As control, no menadione was added to an additional cell dish. Subsequently, the cells were incubated with cycloheximide (100 µg/mL, 5 min). Afterwards, the medium was removed and 10 mL ice-cold PBS supplemented with cycloheximide (100 µg/mL) was added. Cells were harvested by scraping on ice and collected by centrifugation (500 g, 10 min, 4 °C). Cell pellets were stored at -80 °C until further processing.

For the polysome isolation, the cells were resuspended in lysis buffer (5 mM Tris-HCl (pH 7.5), 2.5 mM MgCl_2_, 1.5 mM KCl, complete EDTA-free protease inhibitor cocktail (Roche), 100 µg/mL cycloheximide, 2 mM DTT, 50 U/mL RNasin (Promega)) and incubated on ice for 20 min. The cell suspension was vortexed every 5 min. Subsequently, the lysate was cleared by centrifugation (16.000 g, 5 min, 4°C). For each sample, the absorption at 260 nm was determined and the lysate concentration was adjusted to A_260_ = 20 with lysis buffer for each sample. 500 µL of the lysate was loaded on the pre-prepared sucrose gradients and ultra-centrifuged (36’000 rpm, 2 h, 4°C) using a SW41Ti rotor (Optima L-90K Ultracentrifuge, Beckman Coulter).

For the recording of the polysome profiles, a Piston Gradient Fractionator (Biocomp) was used. The profiles were recorded by measuring the absorption at 260 nm from the top to the bottom of the gradient. Fractions were collected automatically in a volume of 800 µL. Subsequently, 600 µL of the fraction was mixed with 600 µL ice-cold ethanol, snap-freezed in liquid nitrogen and stored at -80 °C.

#### Protein preparation for western blot analysis

The samples containing 50% ethanol were centrifuged (20’000 g, 30 min, 4°C), the supernatant was discarded and the protein pellet was washed with 80% ethanol (500 µL). The pellet was dried at 95 °C for 3 min and the proteins were resuspended in 1x Laemmli sample buffer (fraction 1, 2: 250 µL; fraction 3-12: 25 µL). The samples were denatured (95°C, 5 min) and analyzed by SDS-PAGE and western blot as described above.

#### Puromycin incorporation assay

One day prior to the experiment, 1.2 × 10⁶ HEK293 WT or Rlig1-KO cells were seeded in 6-well plates and incubated as described in the cell culture section above. For oxidative stress induction, cells were treated with 40 µM menadione for 1 h. For the final 15 min of treatment, 5 µM puromycin was added to the well to label newly synthesized polypeptides. Control cells were incubated with puromycin alone. Following the treatment, cells were harvested, and puromycin incorporation was visualized by SDS-PAGE and immunoblotting as described above. Quantification of protein synthesis rate was achieved by calculating the ratio of puromycin signal intensity to total protein levels which were determined by a separate Coomassie staining. Protein synthesis levels in untreated HEK293 WT cells were defined as 100%, and all other conditions were normalized relative to this reference.

## RESULTS

### Work-flow for the identification of Rlig1 interaction partners

In order to identify cellular protein-protein interaction partners of Rlig1 in cell lysates from HEK293 Rlig1-KO cells, we conducted an affinity enrichment coupled to mass spectrometry (Figure 1). We employed recombinantly expressed AMP-Rlig1 as the active bait and included the catalytically inactive K57A-Rlig1 variant, which cannot undergo spontaneous AMPylation during affinity enrichment, to distinguish activity-dependent from activity-independent interactions.(17) An additional uncharged bead control was included to differentiate between non-specific binding to the beads and specific interactions of Rlig1. Cell lysates from HEK293 Rlig1-KO cells were used to minimise competition between endogenous Rlig1 and the employed bait. In line with previous experiments, cell lysates from either unstressed or menadione-stressed cells were used to investigate potential stress-induced changes in the interaction partners of Rlig1.(17) Also, the cell lysates were previously cleared from RNA to exclude indirect RNA-mediated interactors. Here, RNA digestion was optimised with the RNA endonuclease RNase I_f_, as it is a common and well-characterised RNA endonuclease that cleaves RNA dinucleotide bonds without sequence specificity.(34) RNase concentration was optimised for a short incubation time (Figures S1A and B and S2A). The Rlig1 variants were immobilised on streptavidin conjugated agarose beads via a co-expressed Strep-tag and the loaded beads were subsequently incubated with the cell lysates cleared from RNA. After thorough washing of the beads, the enriched proteins were eluted with biotin and prepared for subsequent analysis by SDS-PAGE and MS-based proteomics.

**Figure 1:**
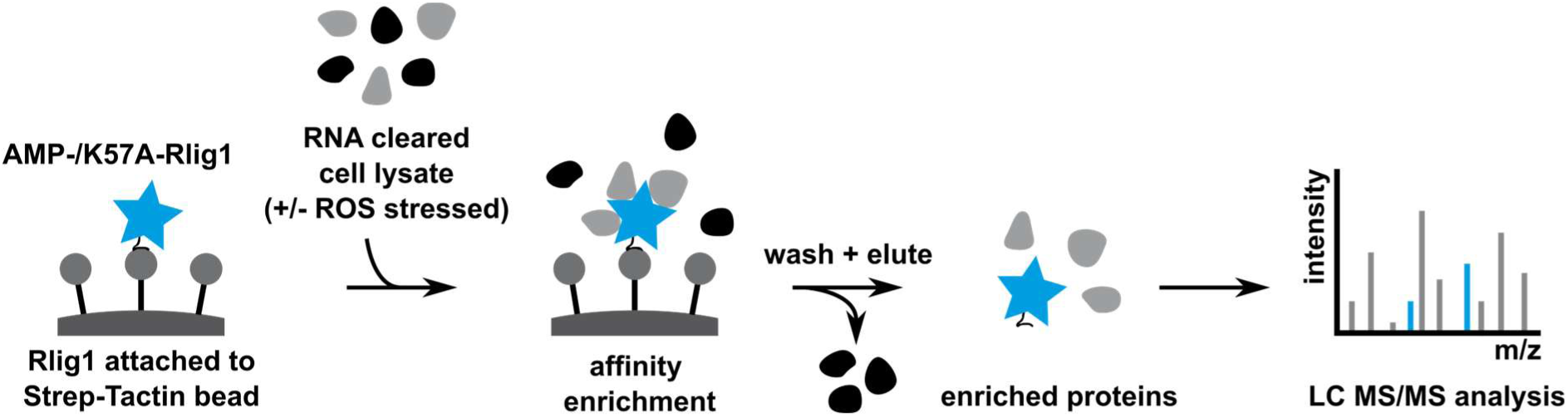
Identification of protein-protein interaction partners of Rlig1. Recombinantly expressed AMP-Rlig1 (enzymatically active form) or K57A-Rlig1 (inactive form) was attached via a co-expressed Strep-tag to Strep-Tactin beads. HEK293 Rlig1-KO cells were grown under physiological conditions (unstressed) or treated with menadione (stressed), lysed and cleared from RNA using RNase I_f_. The RNA cleared cell lysate was incubated with the bait attached to the Strep-Tactin beads. After thorough washing of the beads, potential protein interactors were eluted, digested and identified by SDS-PAGE and mass spectrometry-based proteomics.

### Identification of cellular interaction partners of Rlig1

Eluted proteins were first analysed by SDS-PAGE and visualised by Krypton staining. Here, Rlig1-containing samples showed multiple additional protein bands in comparison to the empty bead control, clearly indicating a successful enrichment (Figure S2B). Furthermore, differences in the intensities of various bands in the enriched proteins of the stressed and unstressed approach could be detected in the gel. Eluted proteins were then subjected to a tryptic in-solution digest and analysed by MS based proteomics using label-free data-independent acquisition (DIA) for identification and quantification of potential Rlig1 interactors. Initial principal component analysis (PCA) of all samples was able to effectively distinguish the bead control from the bait-containing samples in the first principal component (Figure S3). Also, active AMP-Rlig1 samples were clearly separated from inactive K57A-Rlig1 samples. The stressed samples were effectively separated from the unstressed samples in the second principal component. The variance of the biological replicates proved to be significantly smaller than between the different conditions. 2, 107 proteins were confidently identified for the unstressed and 2, 125 for the stressed samples (q-value < 0.01 and identified in all three biological replicates with ≥3 peptides in at least one of the conditions; for details see materials and methods). From those, 588 proteins were significantly enriched by AMP-Rlig1 in the unstressed and 459 in the stressed samples (ANOVA FDR = 0.01, S_0_ = 0.1; Post hoc Tukey’s test FDR = 0.01) (Figure 2A) (for details, see experimental section, for the resulting protein lists see SI). The majority of proteins was enriched for both variants, albeit to a lesser extent for the inactive K57A-Rlig1 in most cases (Figure S4). A large overlap of 384 proteins existed between the stressed and the unstressed samples (84% and 65% of the enriched proteins, respectively). Only 75 proteins were enriched exclusively in the stressed samples, while 204 proteins were solely enriched in the unstressed samples for AMP-Rlig1. Subsequent analysis of the AMP-Rlig1 enriched proteins revealed very similar significant GO terms for the proteins enriched in the two treatments (Figure 2B), as well as for the proteins enriched exclusively in the stressed or unstressed samples (For the lists of GO terms for the different treatments see SI). Overall, our data therefore shows a consistent set of cellular interaction partners for Rlig1 after menadione treatment.

**Figure 2:**
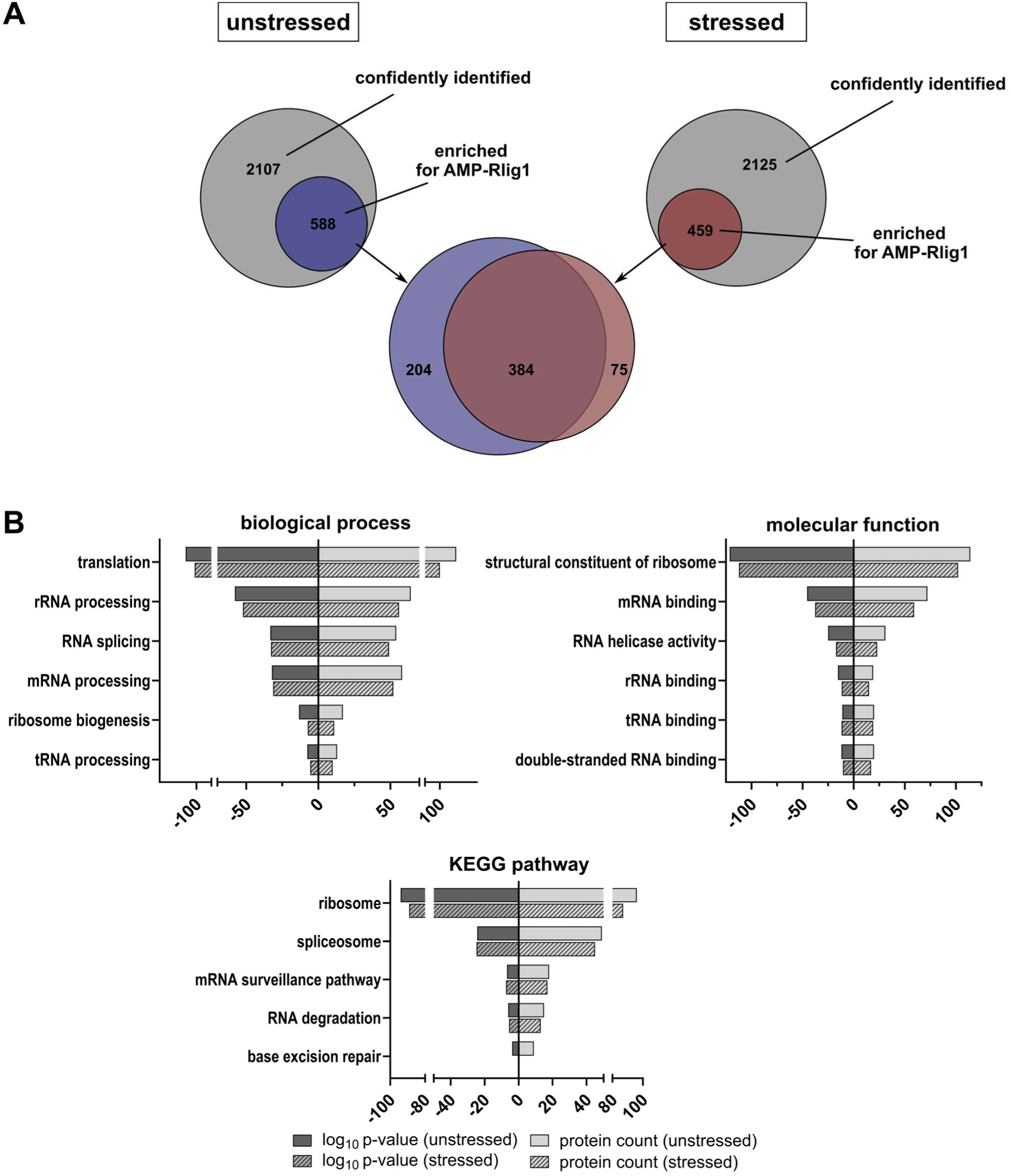
AMP-Rlig1 interaction partners are mainly involved in RNA-based processes and are highly similar for stressed and unstressed cells. A) Venn diagram of identified and significantly enriched proteins for the active AMP-Rlig1 variant from HEK293 Rlig1-KO cells grown under physiological conditions (blue) or after treatment with menadione (red). Significant enrichment determined from biological triplicates by ANOVA multiple sample test (S_0_ = 0.1, FDR = 0.01) and Post hoc Tukey’s test (FDR = 0.01). B) Classification of potential Rlig1 interaction partners according to GO-term analysis. Shown are enriched biological processes, molecular functions and KEGG pathways based on DAVID classification.(29) Plotted are the log_10_ of the modified Fisher Exact p-value for the corresponding GO term and the count of proteins belonging to this GO term. Results for proteins enriched in unstressed and stressed samples were plotted separately as indicated by the legend.

Strongly enriched by AMP-Rlig1 in the stressed and unstressed condition were proteins involved in the biological processes of translation, RNA splicing and the processing of various RNAs. In terms of molecular function, binders of various types of RNA and proteins with RNA helicase activity were noticeably enriched. Here, most of the identified RNA helicases were stronger enriched for the active AMP-Rlig1 variant, than for the inactive K57A-Rlig1 variant (Figure S5A). In regard to significant KEGG pathways, “mRNA surveillance pathway”, “RNA degradation” and in the unstressed cells “base excision repair” did stand in line with a potential role for Rlig1 in RNA damage response.

The majority of potential Rlig1 interactors from the group of “RNA degradation” were members of the exosome. Here, all but one of the known exosome proteins were enriched in both the stressed and the unstressed treatment, strongly suggesting an interaction of Rlig1 with the exosome complex (Figure S5B). Remarkably, DIS3, the catalytically active subunit of the exosome complex, (35) completely changed its binding-preference upon stress treatment. In the unstressed sample, DIS3 was strongly enriched for the enzymatically active AMP-Rlig1 bait. Conversely, DIS3 was almost exclusively enriched for the inactive K57A-Rlig1 in the stressed samples (Figure S5B). The shift in enrichment from the active AMP-Rlig1 variant under unstressed conditions to the inactive K57A-Rlig1 variant under oxidative stress was a rare observation in our dataset and, among the proteins analysed in detail, represented the only instance of such a clear change in binding preference.

Within the group “base excision repair”, various members of both, the short and long-patch base excision DNA repair system were found to be enriched by Rlig1. These involved for example the glycosylase MPG, and the endonuclease APE1 which are both already known to be able to process RNA in addition to their role in DNA repair.(27, 36–38)

Interestingly, two potential interactors of Rlig1 in a hypothetical heal-and-seal RNA cascade, ANGEL2 and CLP1, were significantly enriched for active AMP-Rlig1 in both, stressed and unstressed samples (Figure S5C). These two enzymes could potentially modify RNA strands to enable a subsequent ligation by Rlig1. It was already shown that ANGEL2 and Rlig1 are able to enzymatically cooperate to ligate RNA strands in vitro, further corroborating a role of Rlig1 in RNA repair.(17)

Furthermore, a significant enrichment of ribosomal proteins was observed in the affinity enrichment and the GO term analysis, strongly suggesting an interaction between Rlig1 and the ribosome. Most of these ribosomal proteins showed stronger enrichment for the active AMP-Rlig1 variant than the inactive K57A-Rlig1 variant, in both unstressed and stressed cells (Figure S5D).

### Rlig1 interacts with the ribosome **in vitro**

As our affinity enrichment mass spectrometry data suggested an interaction of Rlig1 with the ribosomal machinery, we sought to validate the interaction between Rlig1 and the ribosome. We employed a ribosome sedimentation experiment, in which different concentrations of purified and intact human 80S ribosome were incubated with recombinantly expressed AMP-Rlig1 and sedimented through a sucrose cushion. The intact 80S ribosome has sufficient mass to traverse the sucrose cushion, whereas AMP-Rlig1 alone is not able to sediment. When AMP-Rlig1 and the ribosome were incubated prior to sedimentation, AMP-Rlig1 was identified in the sediment (Figure 3A). The amount of detected AMP-Rlig1 also correlated with the amount of 80S ribosome used in this assay. However, the RNA ligase did not sediment in an equimolar ratio with the ribosome and the signal for Rlig1 from the sedimented samples was markedly reduced in comparison to the signal before sedimentation. Therefore, the blots showing the Rlig1 signal before and after sedimentation, had to be imaged separately to avoid overshadowing of the weaker signals after sedimentation. Taken together, the results of this assay indicate an interaction of AMP-Rlig1 with the ribosome that is of sufficient strength to survive sedimentation through the sucrose cushion.

**Figure 3:**
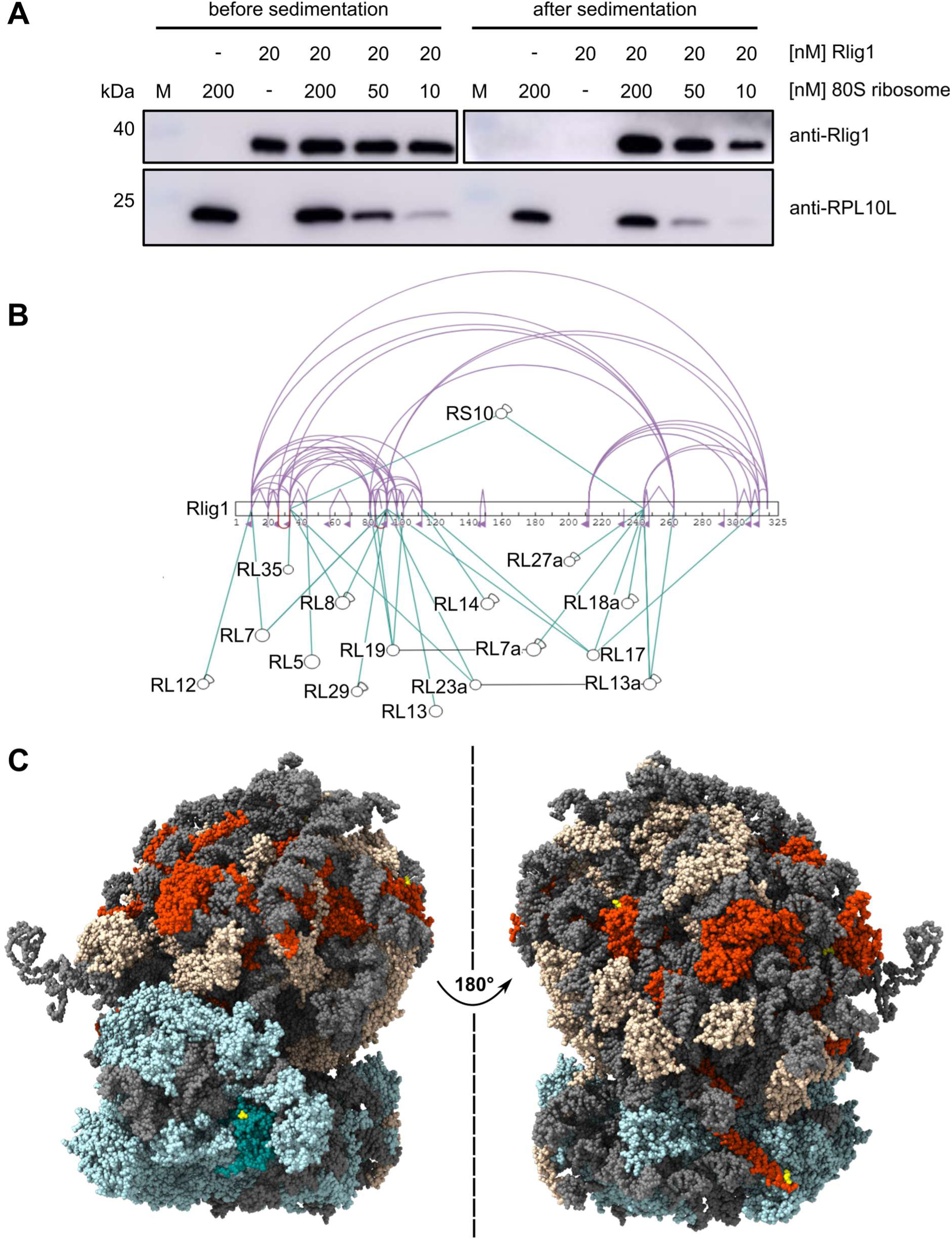
AMP-Rlig1 binds preferentially to the large ribosomal subunit of the 80S ribosome in vitro A) Co-sedimentation of AMP-Rlig1 with different concentrations of human 80S ribosome analysed by western blot. Anti-Rlig1 (top) and anti-RPL10L were used as control for the sedimentation of the 80S ribosome (bottom). Of note: the Rlig1 western blot after sedimentation was exposed for a longer time compared to the one before the sedimentation in order to result in a sufficient signal. B) Crosslinking of AMP-Rlig1 with the 80S ribosome using the lysine reactive crosslinker BS3. Shown are only crosslinks between Rlig1 and ribosomal subunits that were confidently identified in 2 of 3 biologically independent replicates (32). Rlig1 is shown in an opened sequence view while ribosomal proteins are indicated by black circles with arches indicating identified intra-protein crosslinks for this protein. Inter-protein crosslinks of ribosomal proteins to AMP-Rlig1 are shown in turquoise and intra-protein crosslinks of AMP-Rlig1 in violet. Inter-protein crosslinks between ribosomal proteins are shown in black. C) Cryo-EM map of the human 80S ribosome (PDB: 4UG0)(39) showing proteins and interaction sites crosslinked to Rlig1. 60S ribosomal proteins for which a crosslink to Rlig1 was identified are shown in dark orange and the crosslinked small subunit RS10 in teal. Lysines that were directly crosslinked to AMP-Rlig1 are depicted in yellow. 60S ribosomal proteins that were not crosslinked to AMP-Rlig1 are shown in beige, 40S ribosomal proteins that were not crosslinked to AMP-Rlig1 in light blue. The ribosomal RNA is depicted in grey.

#### Rlig1 interacts with the large ribosomal subunit in vitro

After confirming an association of Rlig1 to the ribosome, we tested whether Rlig1 interacts with a specific part of the ribosome. For this, an in vitro crosslinking experiment with purified human 80S ribosome and recombinantly expressed AMP-Rlig1 was conducted. Here the intact ribosome was pre-incubated with AMP-Rlig1 and crosslinked with the homobifunctional, amine reactive crosslinker BS3 (bissulfosuccinimidyl suberate). After digestion by trypsin, the resulting crosslinked peptides were analysed by MS/MS to identify distinct interaction sites between the ribosome and the RNA ligase.

We noticed that the identified inter-protein crosslinks between Rlig1 and the ribosome showed a strong preference for members of the large subunit, as only one of the 16 crosslinked ribosomal proteins (i.e. RS10), was a member of the small ribosomal subunit (Figure 3B).

However, examination of the remaining 15 proteins of the large ribosomal subunit, for which crosslinks were detected, did not reveal a discrete interaction site for AMP-Rlig1 on the large ribosomal subunit. The only obvious common denominator appears to be that the ribosomal proteins are in close vicinity to rRNA on the ribosomal surface (Figure 3C), if mapped onto the cryo-electron microscopy-derived structure of the ribosome (PDB: 4UG0).(39)

### Rlig1 protects translational activity in cells under oxidative stress induced by menadione

After we could further substantiate the binding of Rlig1 to the ribosome in vitro, we investigated the potential role of this interaction and the effect of Rlig1 on ribosomal function in cellulo. We therefore analysed the translation activity of HEK293 WT and HEK293 Rlig1-KO cells by polysome profiling, both under physiological conditions and after treatment with the ROS inducer menadione (Figure 4A). Under physiological conditions, both cell lines showed comparable polysome profiles, characterized by a high distribution of polysomes (Figure 4B). Remarkably, treatment with 40 µM menadione, resulted in a loss of polysomes accompanied by a shift towards the peak for the 80S monosome and those of single ribosomal subunits in Rlig1-KO cells, indicating a decreased translation initiation and translational activity. In contrast, this effect was less pronounced in HEK293 WT cells, where polysome integrity was largely preserved. (Figure 4B, Figure S6A). Treatment with 60 µM menadione resulted in a loss of polysomes and accumulation of 80S ribosomes and the individual subunits in both cell lines (Figure S6A). Still, the peak of the 80S ribosome was more prominent in Rlig1-KO cells than In WT cells, indicating that less polysomes were present in the Rlig1-KO cells (Figure 4B).

**Figure 4:**
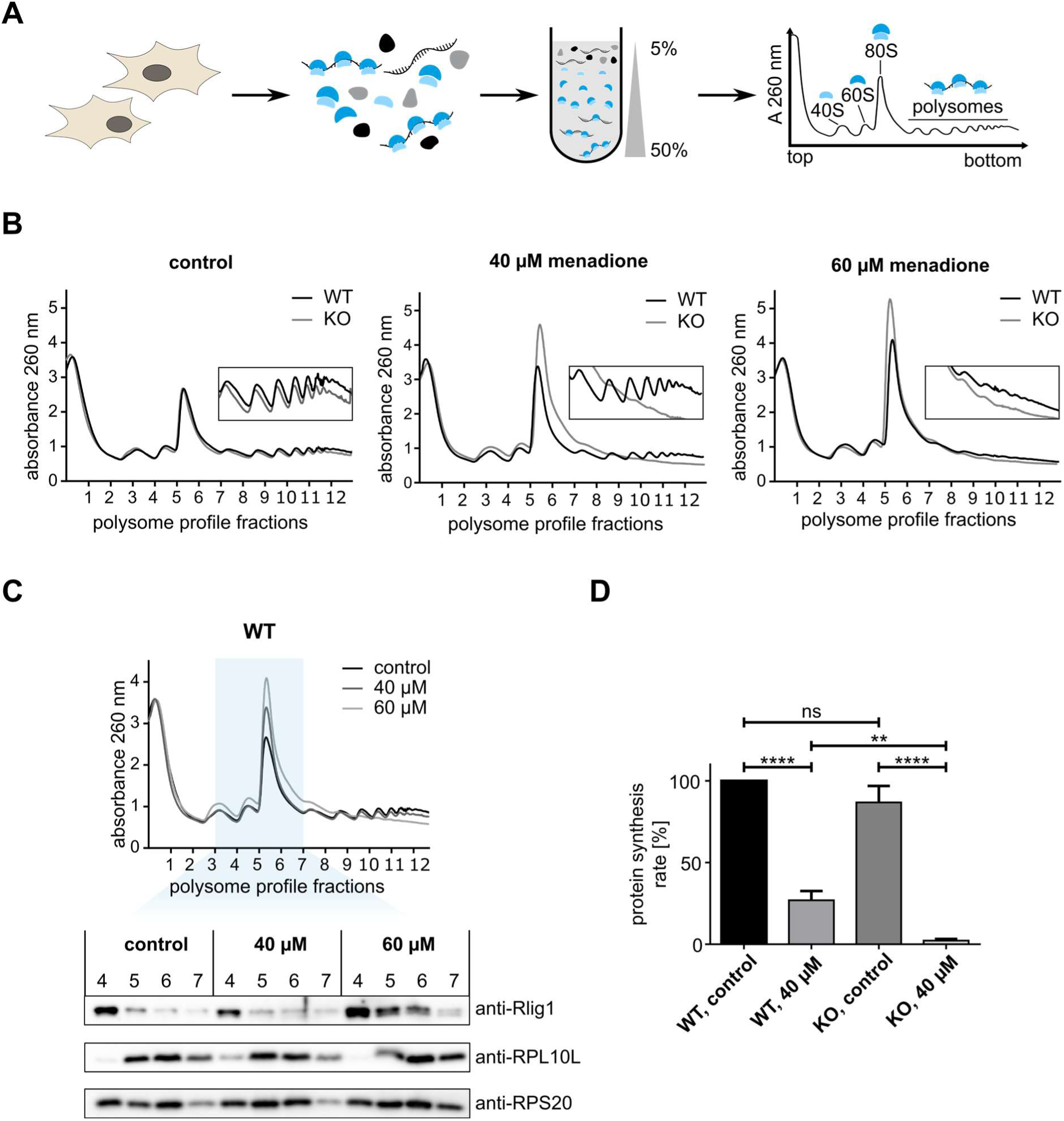
Rlig1 is important for maintenance of translational activity in cells under oxidative stress induced by menadione. (A) Schematic depiction of the polysome profiling workflow. (B) Polysome profile of HEK293 WT (black) and Rlig1-KO cells (grey). Cells were grown under physiological conditions (control) or subjected to 40 µM or 60 µM menadione for 60 min at 37 °C. Translation was arrested with cycloheximide. Cleared cell lysate was separated in a 5–50% sucrose gradient by ultracentrifugation and analysed at 260 nm. (C) Combined polysome profiles of the WT cells treated with different concentrations of menadione (top). Corresponding Western blot analysis of fractions 4 to 7 of the polysome profiles (bottom). Fractions analysed by western blot highlighted in blue. Antibodies were used against RPL10L as control for the 60S ribosomal subunit, RPS20 as control for the 40S ribosomal subunit and against Rlig1. Two independent experiments were performed with similar results. (D) Analysis of protein synthesis rate determined by puromycin incorporation assay in HEK293 WT or Rlig1-KO cells under physiological condition or after treatment with 40 µM menadione for 60 min. Protein synthesis rate was calculated by the ratio of puromycin signal intensity to total protein levels as determined by Coomassie staining. Synthesis rate of untreated HEK293 WT cells was set to 100% and all other values were calculated accordingly. Samples were measured in biological triplicates. Error bars represent the ± SD.

Subsequent immunoblot analysis of the fractions from the polysome profiling of HEK293 WT cells allowed to visualise the relative distribution of Rlig1 in the different fractions. In all conditions, the by far strongest Rlig1 signal could be detected in the soluble, non-ribosomal fractions 1 to 3 (Figure S6B). Due to the high signal intensity in those fractions, the other fractions were analysed separately. To enable a direct quantitative comparison of Rlig1 levels between the different treatment conditions, fractions 4 to 7 of each condition were analysed on the same gel (Figure 4C). These fractions represent the two ribosomal subunits (40S and 60S ribosomal subunit), the 80S monosome, and the initial polysome peak. However, due to automated fraction collection, individual fractions may span multiple adjacent peaks of the polysome profile, preventing a perfect assignment to specific ribosomal units. Under physiological conditions and after treatment with 40 µM menadione, immunoblot analysis indicated the presence of Rlig1 in fraction 4, mainly representing the small 40S ribosomal subunit. However, upon 60 µM menadione treatment, Rlig1 was also detected in fraction 5 and 6, indicating a recruitment of Rlig1 to the large 60S ribosomal subunit and the 80S monosome. This recruitment coincided with the observed loss of polysomes in HEK293 WT cells under these stress conditions.

To further elucidate the role of Rlig1 on cellular translation, global protein synthesis was assessed in HEK293 WT and HEK293 Rlig1-KO cells under physiological conditions and under oxidative stress induced by 40 µM menadione. Puromycin labelling was conducted to assess translational activity in the cells (Figure S6C). Under physiological conditions, no significant difference in global protein synthesis was observed between HEK293 WT and Rlig1-KO cells (Figure 4D). Upon oxidative stress however, both cell lines exhibited a marked reduction in protein synthesis. Notably, this decrease was significantly more pronounced in Rlig1-KO cells compared to the WT cells. Those findings, together with the observed accumulation of 80S monosomes in Rlig1-KO cells under oxidative stress and the stress-dependent redistribution of Rlig1 to ribosome-associated fractions, suggest that Rlig1 plays a role in maintaining translational activity under the stress conditions induced by menadione.

## DISCUSSION

In recent years, the possibility of RNA repair in response to damaged RNAs as an alternative to RNA degradation processes has come into focus. However, besides the reversal of specific methylations(6) and the resolution of RNA-protein crosslinks, (7–9) no additional RNA repair pathways have been identified to date. In a previous study, we identified Rlig1 as the first human RNA ligase to employ a 5’-3’ ligation mechanism, (17) which indicated the possible existence of yet unidentified RNA repair mechanisms in human cells.

Although the enzymatic activity of Rlig1 has been previously characterised, a potential involvement of Rlig1 as an RNA ligase in specific cellular pathways remains to be elucidated.(17, 20–22) Here, we report the successful identification of potential cellular interaction partners of Rlig1 by affinity enrichment mass spectrometry. Subsequent experiments indicate a possible role for Rlig1 in maintaining mature ribosomes and in maintaining translation upon oxidative stress induced by menadione. Taken together, we provide further evidence for a functional role of Rlig1 as an RNA ligase relevant for cellular fitness under stress conditions.

Among the identified possible interactors of Rlig1 are proteins involved in the biological processes of translation, processing of RNAs and splicing. Furthermore, proteins involved in “mRNA surveillance”, “RNA degradation” and “base excision repair” were found to be significant enriched, strengthening the idea of Rlig1 being important for RNA maintenance or repair.

Among the enriched proteins involved in RNA degradation were predominantly members of the exosome. Interestingly, the affinity of DIS3, the catalytic subunit of the exosome complex, (35) for AMP-Rlig1 was oxidative stress dependent. Under physiological conditions, DIS3 was strongly enriched for active AMP-Rlig1. However, under oxidative stress induced by menadione, DIS3 was no longer enriched for AMP-Rlig1. These results indicate that Rlig1 interacts with DIS3 and the exosome in a stress-sensitive manner, suggesting a possible involvement of Rlig1 in exosome-associated RNA surveillance pathways.

Also, the enrichment of proteins involved in the base excision repair of DNA is intriguing. In particular, as some of the identified potential interactors of Rlig1 in this study are known to be able to process also RNA.(25–27) Known examples for such RNA processing base excision repair proteins that were found to be enriched by Rlig1 are the glycosylase MPG1, (27, 36) and the endonuclease APE1, (27, 37, 38) as well as its interaction partner NPM1 which regulates the activity of APE1 on RNA substrates depending on its binding status.(40)

A possible involvement of Rlig1 in a hypothetical RNA repair pathway was further supported by our finding that the two enzymes ANGEL2 and CLP1 were significantly enriched with the active AMP-Rlig1. These two enzymes could, in theory, process RNA strands that result from a classical RNA strand break involving intramolecular transesterification. Such a strand break results in the formation of a 2’, 3’-cyclic phosphate (cPO_4_) group and a 5’-OH group, which are initially incompatible with Rlig1-mediated ligation.(17, 41) In this scenario, CLP1 could add the required phosphorylation to the RNA 5’-end of the RNA strand, and ANGEL2 could remove the 2’, 3’-cPO_4_, preparing the strands for subsequent ligation by Rlig1.(42, 43) TRPT1 and CNPase, which could potentially act as surrogates for ANGEL2 in such a scenario, were not enriched in our data.(44, 45) ANGEL2 and Rlig1 have previously been shown to be able to enzymatically cooperate in the ligation of RNA strands in vitro.(17) The identification of these proteins as interactors of Rlig1 further supports the concept that these proteins may cooperate with Rlig1 in order to repair damaged RNA.

A large proportion of the proteins that were found to be enriched for the active AMP-Rlig1 were also enriched for the inactive K57A-Rlig1 variant, although often to a lesser extent. The AMPylation state of Rlig1 seems therefore to mostly impact the strength of a particular protein-protein interactions of Rlig1 but does not appear to change the interactors per se to a great extent. Similarly, the applied stress treatment had little impact on the overall molecular functions of Rlig1 interaction partners. Most enriched proteins were shared between the unstressed and stressed conditions, and enriched GO terms were largely overlapping. Even proteins that were uniquely identified in either condition showed similar functional annotations. These findings suggest that the Rig1-associated protein network remains largely stable under stress. This could hint at a role for Rlig1 in general RNA maintenance beyond stress conditions.

Interestingly, ribosomal proteins were found to extensively bind to Rlig1. Subsequent experiments comprising a ribosome sedimentation as well as an in vitro ribosome crosslinking experiment using recombinant AMP-Rlig1 confirmed this interaction of AMP-Rlig1 with the ribosome. However, in the ribosome sedimentation experiment, Rlig1 did not co-sediment in a 1:1 ratio with the ribosome. This suggests, that either the binding might not be strong enough to guarantee the complete sedimentation or that AMP-Rlig1 only binds to the ribosome under specific conditions. Our in vitro crosslinking experiment confirmed binding of Rlig1 to the ribosome, preferentially to its large subunit, even though a specific interaction area could not be identified. Rather, our data indicated an association with accessible RNA patches on the ribosomal surface, corroborating a potential role of Rlig1 in RNA repair. This finding was intriguing, as earlier research had already demonstrated that Rlig1 is crucial in delaying the degradation of the 28S rRNA of the large ribosomal subunit upon oxidative stress with menadione.(17)

To investigate whether the in vitro interaction of AMP-Rlig1 with the ribosome has functional relevance in a cellular context, we examined the impact of Rlig1 on translational activity utilising polysome profiling and a puromycin incorporation assay. These analyses revealed that oxidative stress impairs translation more severely in the absence of Rlig1. In Rlig1-KO cells, polysome loss accompanied by accumulation of the 80S ribosome and its subunits occurred at lower menadione concentrations compared to WT cells, indicating increased translational sensitivity to oxidative stress. This effect was further corroborated by a marked decrease in global protein synthesis as evidenced by the puromycin assay. In contrast, WT cells maintained a higher translational activity under oxidative stress induced by menadione, suggesting some kind of safeguarding role of Rlig1 for the translation. Under the stress conditions induced by treatment with 60 µM menadione however, polysomes were also strongly reduced in WT cells, which coincided with the observed stress-induced recruitment of Rlig1 to ribosomal subunits and the 80S monosome.

In light of the data presented in this study, our results suggest that the interaction of Rlig1 with the ribosome, and its increased association under menadione-induced oxidative stress, may be important for maintaining polysomes and cellular translational activity. The affinity of Rlig1 for proteins with the potential to repair RNA strand breaks, as well as for DNA repair factors with reported enzymatic activity on RNA, is consistent with the possibility of a yet unidentified RNA repair pathway involving Rlig1. Further work will be required to clarify how the interaction between Rlig1 and the ribosome is regulated, and whether Rlig1 can act in cooperation with other enzymes, such as ANGEL2 and CLP1, to support RNA integrity more broadly and ribosomal RNA in particular.

## Supporting information

Lists of significantly enriched proteins

Lists of enriched GO terms

## AUTHOR CONTRIBUTIONS

Conzeptualization: F.M.S and A.M.; Investigation, data curation, formal analysis and visualization: F.M.S, M.G., J.J.; Resources and supervision: F.S. and A.M.; Project administration and funding acquisition: A.M.; Writing – original draft: F.M.S. and M.G.; Writing- review & editing: F.M.S, M.G., J.J., F.S. and A.M.

## SUPPLEMENTARY DATA

The lists of significant enriched proteins of the affinity enrichment coupled to mass spectrometry and the lists of the corresponding enriched GO terms are included in the supplementary information files.

Raw data of the affinity enrichment coupled to mass spectrometry, as well as the data bases utilised, and the files of the data analysis are available via ProteomeXchange with the identifier PXD067856 (Username; Password: 5cD8925AZ6ug).

Data of the XL-MS experiment are available via ProteomeXchange with the identifier PXD067615 (Username; Password: 7nHY32A9Vzj6).

## CONFLICT OF INTEREST

The authors declare no competing interests.

## FUNDING

A.M. acknowledges the European Research Council for funding of this project (ERC AdG AMP-Alarm, 101019280).

F.S. is grateful for funding by German Research Foundation (DFG: 496470458 and 516836828).

## Notes

### Competing Interest Statement

The authors have declared no competing interest.

### Summary of Updates

In the previous version important supplemental files were missing. The supplemental files were updated.

