## Supplementary Figures for "Human RNA ligase 1 as a novel regulator of ribosome function and translation under oxidative stress"

**MANUSCRIPT TITLE**

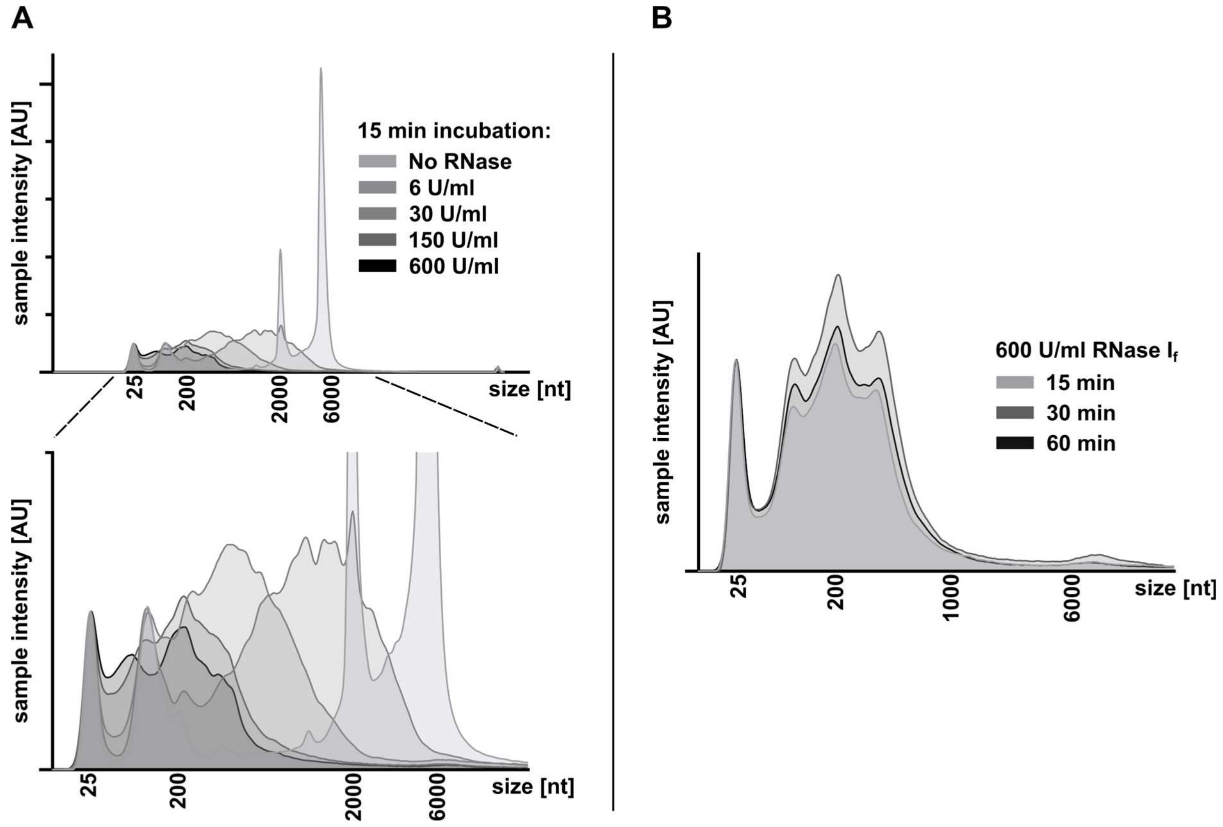


**Figure S1: Establishment of the RNA digestion for the affinity enrichment.** A) Titration of different concentrations of RNase $I_f$ for optimal digestion with 15 min incubation time. The bottom graph is a close-up of the upper graph. B) Determination of the incubation time needed for an optimal RNA degradation with 600 U/mL RNase $I_f$.

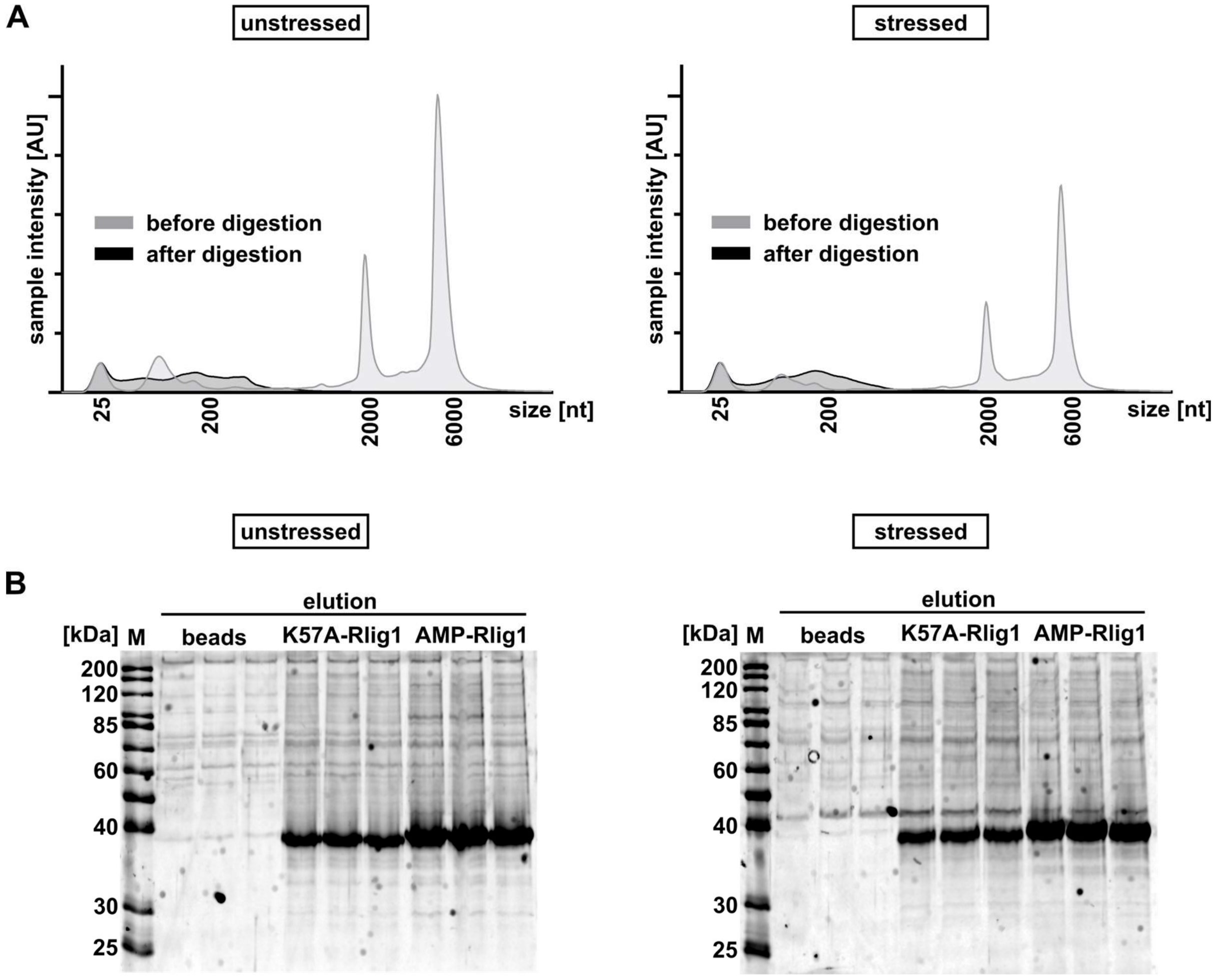


**Figure S2: Control of the RNA digestion prior to the affinity enrichment and Krypton-stained gels of the elution samples.** A) Exemplary TapeStation 4150 analysis of the RNA in the cell lysates directly after cell lysis (light grey) and of the RNase $I_f$ digested cell lysate directly before application of the lysate to the beads (in black). On the left are the analyses for the unstressed samples, on the right for the stressed samples. B) Krypton-stained gels showing the protein content of the eluted protein samples before trypsinisation. On the left the gel of the unstressed samples, on the right of the stressed samples.

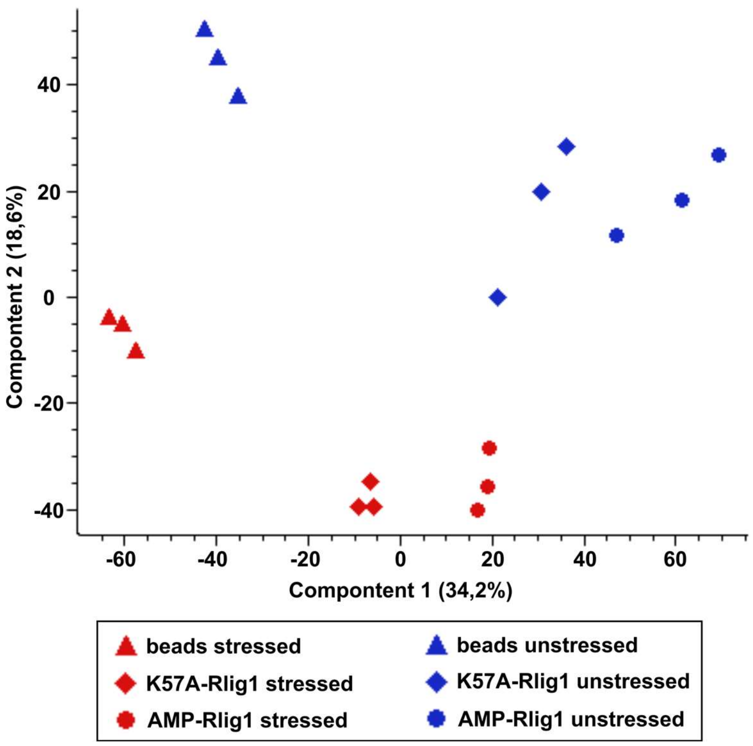


**Figure S3: Principal component analysis (PCA) of the identified proteins of the affinity enrichment samples in triplicates.** On the X-axis component 1, on the Y-axis component 2. Stressed samples in red, unstressed samples in blue.

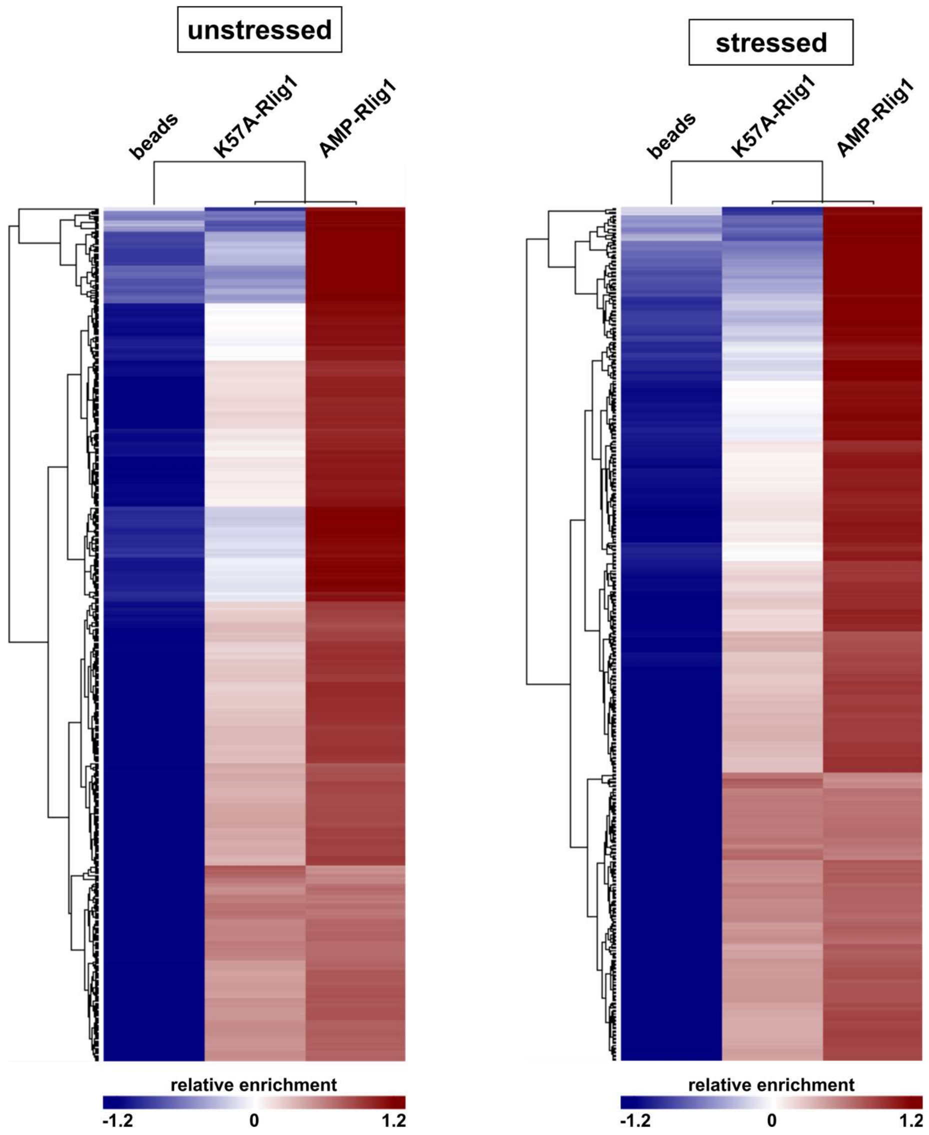


**Figure S4: Heatmap of the significantly enriched proteins for AMP-Rlig1.** Hierarchical clustering (Euclidean distance) of statistically significantly enriched interactors of AMP-Rlig1 following ANOVA analysis (S0 = 0.1, FDR = 0.01) and Post-Hoc Tukey's test (FDR = 0.01). Only proteins with a minimum Z-score of >0.5 for the AMP-Rlig1 bait and a Z-score <0 for the bead control after the ANOVA analysis are shown. Columns represent the sample type, rows represent an interacting protein. On the left side the heatmap for the unstressed enrichment, on the right side for the stressed enrichment. Colouring signals the relative enrichment of the enriched protein as indicated in the legend.

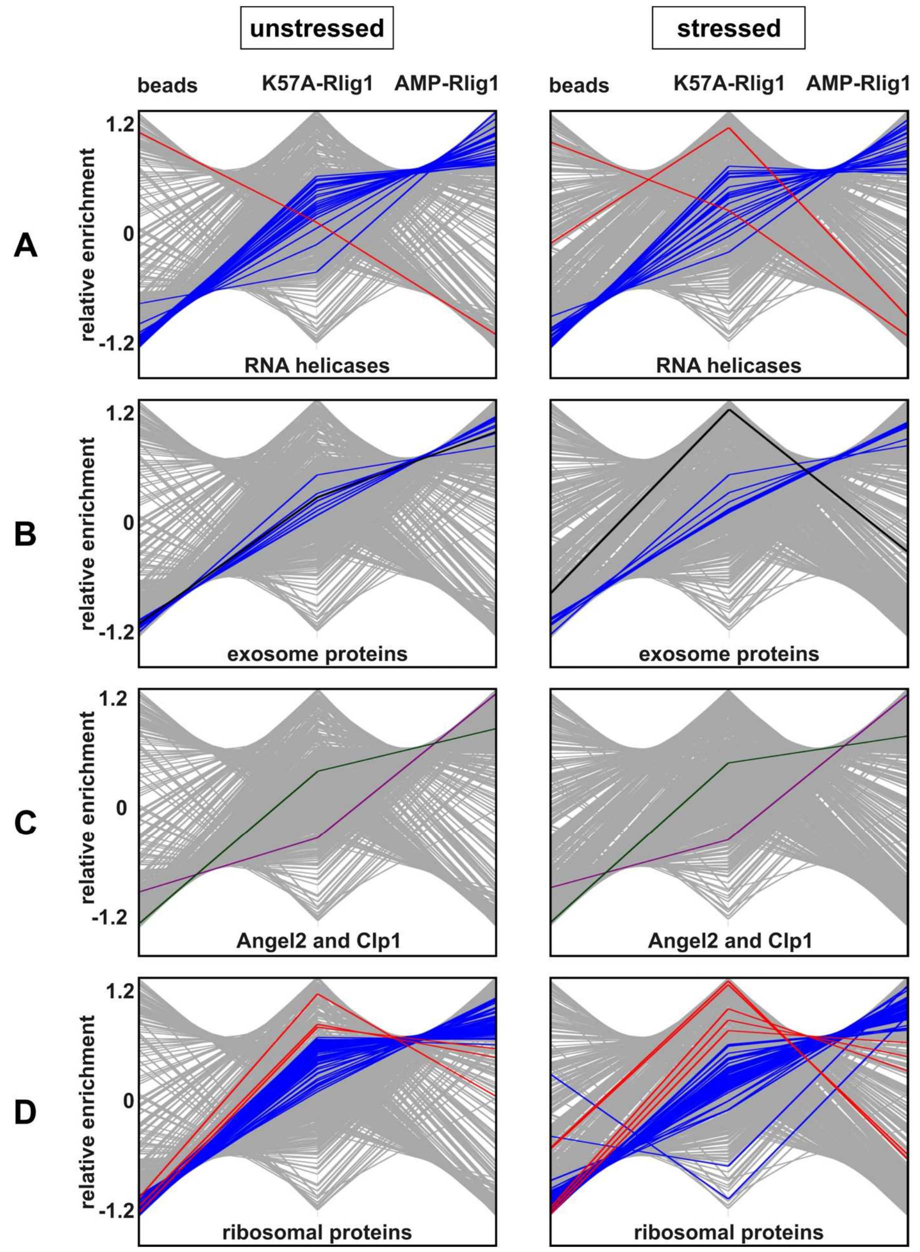


**Figure S5: Profile plots of different Rlig1 interacting proteins.** Proteins belonging to the group of interest are depicted in colour. Proteins enriched the most for AMP-Rlig1 depicted in blue, proteins enriched the most for K57A-Rlig depicted in red. In the background in grey all the other identified proteins. A) RNA helicases (PG.ProteinDescriptions = RNA helicase). B) Proteins of the exosome. The catalytically active exo- and endoribonuclease DIS3/RRP44 is depicted in black. C) The 2′,3′-cyclic phosphatase Angel2 in green, the RNA kinase CLP1 in violet. D) Cytosolic ribosomal proteins of the large (60S) and the small (40S) ribosomal subunit (PG.ProteinDescriptions = 60S/40S).

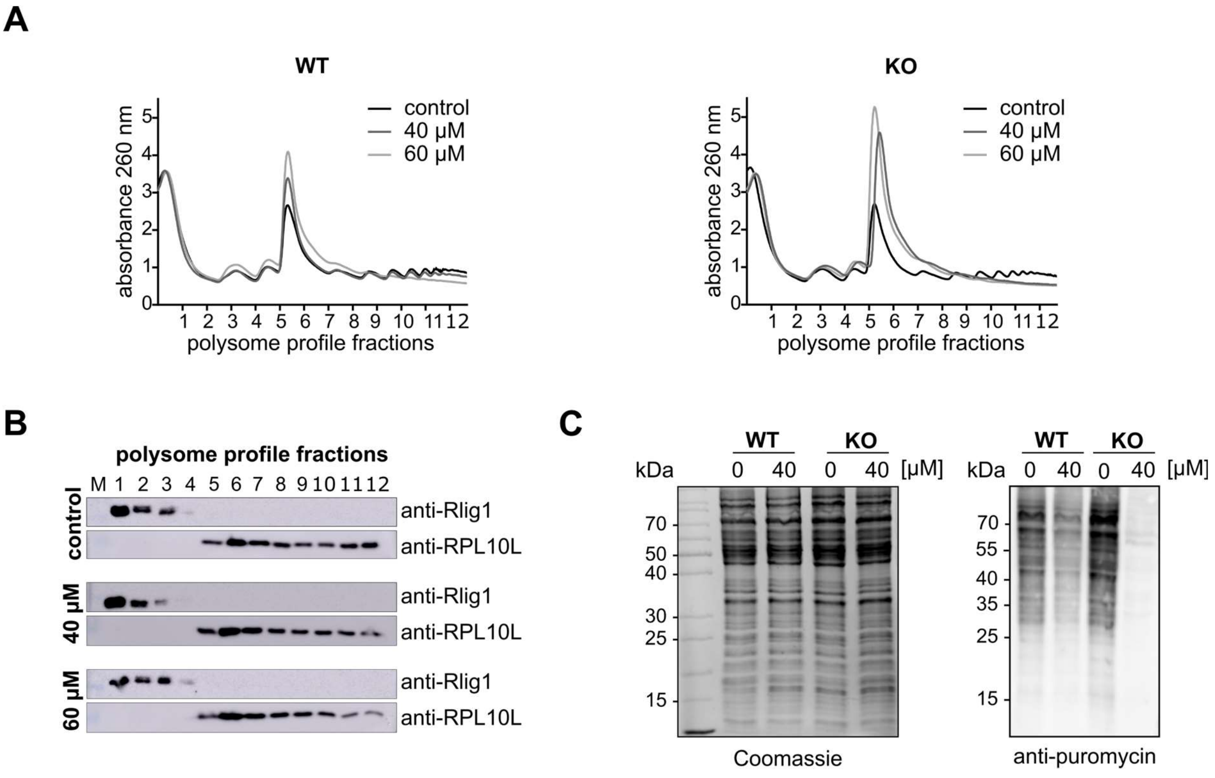


**Figure S6: Analysis of polysome profiles and translational activity in HEK293 WT and Rlig1-KO cells under oxidative stress induced by menadione.** A) Combined polysome profiles of the WT (left) or Rlig1-KO cells (right) treated with increasing concentrations of menadione. B) Western blot analysis of polysome profile fractions of WT cells treated with different concentrations of menadione. Antibodies against RPL10L as control for the 60S ribosomal subunit and against Rlig1 were used. C) Puromycin incorporation assay in WT and Rlig1-KO cells, either untreated or treated with 40 µM menadione. Total protein levels were visualized by SDS-PAGE followed by Coomassie staining (left), and puromycin incorporation was detected by Western blot using an anti-puromycin antibody (right).
